# Postoperative Automated Platelet Count and Its Predictive Factors in Dogs Undergoing Mitral Valve Repair

**DOI:** 10.1101/2025.11.10.687737

**Authors:** Kentaro Kurogochi, Shimon Furusato, Yasuyuki Nii, Ayaka Chen, Yu Ueda, Masashi Mizuno, Masami Uechi

## Abstract

**Objectives:** This study aimed to describe early postoperative changes in automated platelet counts (PLT) after mitral valve repair (MVR) in dogs with myxomatous mitral valve disease (MMVD) and to identify factors associated with these changes.

**Design:** Single-center retrospective cohort study.

**Animals:** Dogs with MMVD that underwent MVR from July 2022 to July 2023. After data extraction, the ten most common breeds were selected, and mixed breeds as well as breeds known for inherent platelet abnormalities (Cavalier King Charles Spaniels and Norfolk Terriers) were excluded.

**Measurements and Main Results:** A total of 313 dogs representing seven breeds met the inclusion criteria. PLT levels were assessed preoperatively (Day 0) and daily for 4 days postoperatively (Days 1–4). Mean (standard deviation) PLT values were 393±127 ×10³/µL on Day 0, 159±72 on Day 1, 137±67 on Day 2, 157±87 on Day 3, and 187±104 on Day 4. Postoperative thrombocytopenia (<100 ×10³/µL) occurred in 124 of 313 dogs during Days 1–4. In a multivariable mixed-effects model, PLT declined significantly after surgery (Day 0 to 1; β = −207.17 [×10³/µL per day], P < 0.001) and showed a postoperative upward trend (Day 1 to 4; β = 20.62 [×10³/µL per day], P = 0.079). Older age was associated with higher PLT; Toy Poodles and Yorkshire Terriers demonstrated lower PLT than Chihuahuas; dogs with ACVIM Stages C– D had higher PLT compared with Stage B2; lower body weight correlated with lower PLT; and intact dogs tended to have higher PLT than neutered dogs.

**Conclusions:** PLT decreased sharply following MVR and demonstrated partial recovery within 5 days. Age, breed, clinical stage, body weight, and neuter status were associated with postoperative PLT patterns, indicating their importance for perioperative assessment and monitoring.

## Introduction

Myxomatous mitral valve disease (MMVD) is the most common cardiac disease in dogs, primarily affecting small breeds [1]. The prognosis for dogs with MMVD and prior congestive heart failure is poor, with a median survival time of less than 1 year when treated solely with medical management [2]. Mitral valve repair (MVR) is an established intervention for dogs with advanced MMVD [3, 4], and although it can extend survival, perioperative management remains insufficiently defined because of limited data.

In human cardiac surgery, platelet counts (PLTs) typically decrease by 40–50% during the first 72 h postoperatively [5]. Postoperative PLT reductions have been linked to hemostatic blood product administration and intrathoracic hemorrhage [6, 7]. A typical presentation involves two successive episodes of PLT fluctuations: an initial early postoperative PLT decline, followed by partial or complete PLT recovery, and then a subsequent unexpected PLT decline, potentially due to immune-mediated mechanisms [8]. In contrast, postoperative PLT dynamics in dogs undergoing cardiac surgery are poorly described. Based on the authors’ clinical experience, dogs frequently experience an early postoperative PLT decline similar to that observed in human patients. Defining these patterns may enhance understanding of PLT dynamics in cardiac surgery and support improved postoperative monitoring and perioperative management in veterinary clinical settings. This study aimed to describe postoperative PLT changes following MVR and identify factors associated with early postoperative PLT fluctuations.

## Animals, Materials, and Methods

### Animals

This retrospective cohort study evaluated postoperative PLT changes in dogs that underwent MVR. The target population included client-owned dogs with MMVD who were candidates for MVR and without known inherited platelet abnormalities. The source population consisted of dogs from this target group examined or treated at the JASMINE Veterinary Cardiovascular Medical Center in Japan. Eligible dogs were those classified as ACVIM stage B2, C, or D based on preoperative echocardiography, with stages C and D requiring documented congestive heart failure [1], and who underwent MVR between July 2022 and July 2023. After data collection, the ten most common breeds were selected, and mixed-breed dogs and breeds predisposed to inherited platelet abnormalities were excluded. The Institutional Ethics Committee of the JASMINE Veterinary Cardiovascular Medical Center approved the study protocol (approval number: 241029-6), and clinical data were collected only after owners provided consent.

### Materials and Methods

#### Mitral Valve Repair

All surgical procedures were performed by two experienced surgeons (MU and MM). Cardiopulmonary bypass (CPB) was conducted with hypothermic management, in which central cooling reduced body temperature to 28–30 °C [9]. After anesthesia induction, approximately 10 mL/kg of whole blood was collected from the jugular vein, stored with acid citrate–dextrose solution, and later reinfused after CPB weaning as autotransfusion. This routinely employed institutional technique aimed to limit postoperative hemodilution, reduce CPB-associated blood cell injury, and preserve coagulation factors. The decision to use autotransfusion was based on pre-CPB hematocrit and hemodynamic assessment. Before CPB cannulation, heparin sodium (200 U/kg) was administered intravenously, and the activated clotting time was verified to exceed 300 s using an analyzer (Sonoclot; Sienco, Inc., Colorado, USA) with a cuvette (gbACT+ Kit; Sienco, Inc., Colorado, USA). Arterial CPB cannulation was placed in the left carotid artery, and venous cannulation was performed via the left jugular vein into the right atrium [10].

For preparation of the CPB circuit, a pediatric oxygenator (CAPIOX FX05 Oxygenator, TERUMO Corporation, Tokyo, Japan) was used, with a minimum circuit volume of 140 mL. The priming solution consisted of 40 mL of water for injection, 0.5 mL of heparin sodium, 2 mL/kg of 7% sodium bicarbonate, 5 mL/kg of 20% D-mannitol, and acetate Ringer’s solution as needed to reach a total volume of 140 mL or approximately 20 mL/kg, depending on each dog’s body size and hydration status. When the expected hematocrit after CPB priming dilution was below 20%, packed red blood cells were added along with acetate Ringer’s solution. The expected post-dilution hematocrit was estimated using a circulating blood volume of 90 mL/kg. The dilution ratio was calculated by dividing the circulating blood volume by the sum of the circulating blood volume and the CPB priming volume. The expected hematocrit was obtained by multiplying the pre-CPB hematocrit measured under anesthesia by this dilution ratio.

Thoracotomy was performed on the left side of the chest through the fourth or fifth intercostal space, and the pericardium was incised. Aortic cannulation was completed after the fat pad surrounding the aortic root was dissected and retracted, followed by clamping of the area with a vascular clamp [10, 11]. The blood-based multi-dose St. Thomas cardioplegia technique was used to achieve cardiac arrest and myocardial protection; infusion was maintained at 5–10 min intervals after arrest. The solution was prepared by mixing crystalloid St. Thomas solution with circulating CPB blood in a 1:1 ratio and adjusting the final potassium concentration to 40 mEq/L [9, 12]. The volume of cardioplegia solution administered was included in the total crystalloid volume. Intracardiac techniques, including artificial chordae and annuloplasty, followed previously described approaches [10, 13]. Based on surgical records, the authors collected details regarding CPB management, including total CPB time, cross-clamp time, time intubated, total cardioplegia volume, intraoperative autotransfusion use, donor transfusion use, and any need for intraoperative re-clamping for surgical correction. Total CPB time was defined as the duration from initiation to cessation of CPB; cross-clamp time was defined as the duration from aortic cross-clamping to clamp release; and time intubated was defined as the interval from endotracheal intubation to extubation.

Postoperative inpatient antithrombotic therapy was initiated with dalteparin sodium (Sawai Pharmaceutical Co., Ltd., Osaka, Japan) at 25 IU/kg at 6 h and 50 IU/kg at 12 h postoperatively. Beginning the day after surgery, treatment consisted of clopidogrel (Plavix, Sanofi-Aventis, Paris, France) at 3.0 mg/kg orally (PO) every 24 h, along with either dalteparin sodium at 100 IU/kg subcutaneously every 8 h or rivaroxaban (Xarelto, Bayer, Leverkusen, Germany) at 0.5–1.0 mg/kg PO every 24 h during hospitalization. After discharge, clopidogrel at 3.0 mg/kg PO every 24 h was continued for 3 months.

#### Complete Blood Count (CBC)

Automated CBC data were obtained using the Celltac α VET (MEK-6550, Nihon Kohden, Tokyo, Japan). CBC values were evaluated preoperatively on the day of surgery (Day 0) and postoperatively from the day after surgery (Day 1) through Day 4 as part of the routine assessment for all dogs. PLT (×10³/µL) across the observation period were collected as the primary outcome. platelet distribution width (PDW) (%), mean platelet volume (MPV) (fL), white blood cell count (/µL), and differential counts for neutrophils (/µL), lymphocytes (/µL), monocytes (/µL), and eosinophils (/µL) were assessed as baseline variables. For the secondary analysis, postoperative thrombocytopenia was defined as a PLT below 100 (×10³/µL) between Day 1 and Day 4. Changes in PDW (%) and MPV (fL) during the observation period were documented as supplemental analyses.

#### Statistical analysis

PLT values were visualized using a spaghetti plot to depict individual trajectories over time. A mixed effects model was generated to evaluate the effects of relevant variables on PLT counts. The time axis was modeled with linear splines and a single knot at Day 1, capturing two phases: perioperative transition (Day 0–1) and postoperative recovery (Day 1–4). Fixed effects included time (day; continuous), breed (selected breeds; polynomial), ACVIM stage (B2, C, D; polynomial), age (years; continuous), body weight (kg; continuous), sex status (neutered male, intact male, neutered female, intact female; polynomial), total CPB time (min; continuous), cardioplegia volume (mL/kg; continuous), intraoperative autotransfusion use (yes, no; binomial), intraoperative donor transfusion use (yes, no; binomial), and need for intraoperative re-clamping (yes, no; binomial). Random effects were assigned to each time period and individual dog. Interaction terms between time period and the covariates were included to assess subgroup differences in temporal trends. A random intercept was applied to each dog and time period to account for repeated measurements and individual variability.

The Shapiro–Wilk test was used to examine the distribution of each continuous variable. Normally distributed variables are reported as the mean ± standard deviation, whereas non-normally distributed variables are shown as the median [interquartile range]. Categorical variables are expressed as counts. The Tukey test was used for group comparisons of normally distributed continuous variables, and the Steel–Dwass test was applied to non-normally distributed variables. Nominal variables were compared using Fisher’s exact test with the Holm– Bonferroni correction. Statistical significance was defined as P < 0.05. All analyses were performed using R software (R version 4.2.2, Foundation for Statistical Computing, Vienna, Austria).

## Results

Of the 422 dogs eligible for surgery during the study period, 398 belonging to the ten most common breeds were included. Eighty-five dogs were excluded because they were mixed breed (n = 64) or belonged to breeds with known abnormalities in PLT values related to macrothrombocytopenia (Cavalier King Charles Spaniel, n = 14; Norfolk Terrier, n = 7) [14, 15]. The final cohort consisted of 313 dogs: Chihuahua (n = 175), Toy Poodle (n = 57), Maltese (n = 36), Pomeranian (n = 22), Shih Tzu (n = 10), Yorkshire Terrier (n = 7), and Miniature Schnauzer (n = 6). Among them, 48 were intact males, 135 were neutered males, 6 were intact females, and 124 were neutered females. Median age was 11.0 years [9.7–12.3 years], and median body weight was 3.3 kg [2.6–4.2 kg]. ACVIM stage distribution was 111 in B2, 163 in C, and 39 in D. Preoperative CBC values showed a PLT count of 392±128 ×10^3^/µL, a white blood cell count of 8,300/µL [6,400–10,200/µL], neutrophils at 5,900/µL [4,400–7,500/µL], lymphocytes at 1,700/µL [1,400–2,300/µL], monocytes at 300/µL [200–400/µL], and eosinophils at 100/µL [0– 100/µL]. PLT, white blood cell count, neutrophils, and monocytes were higher in dogs with more advanced ACVIM stage (**Table 1**).

**Table 1.**
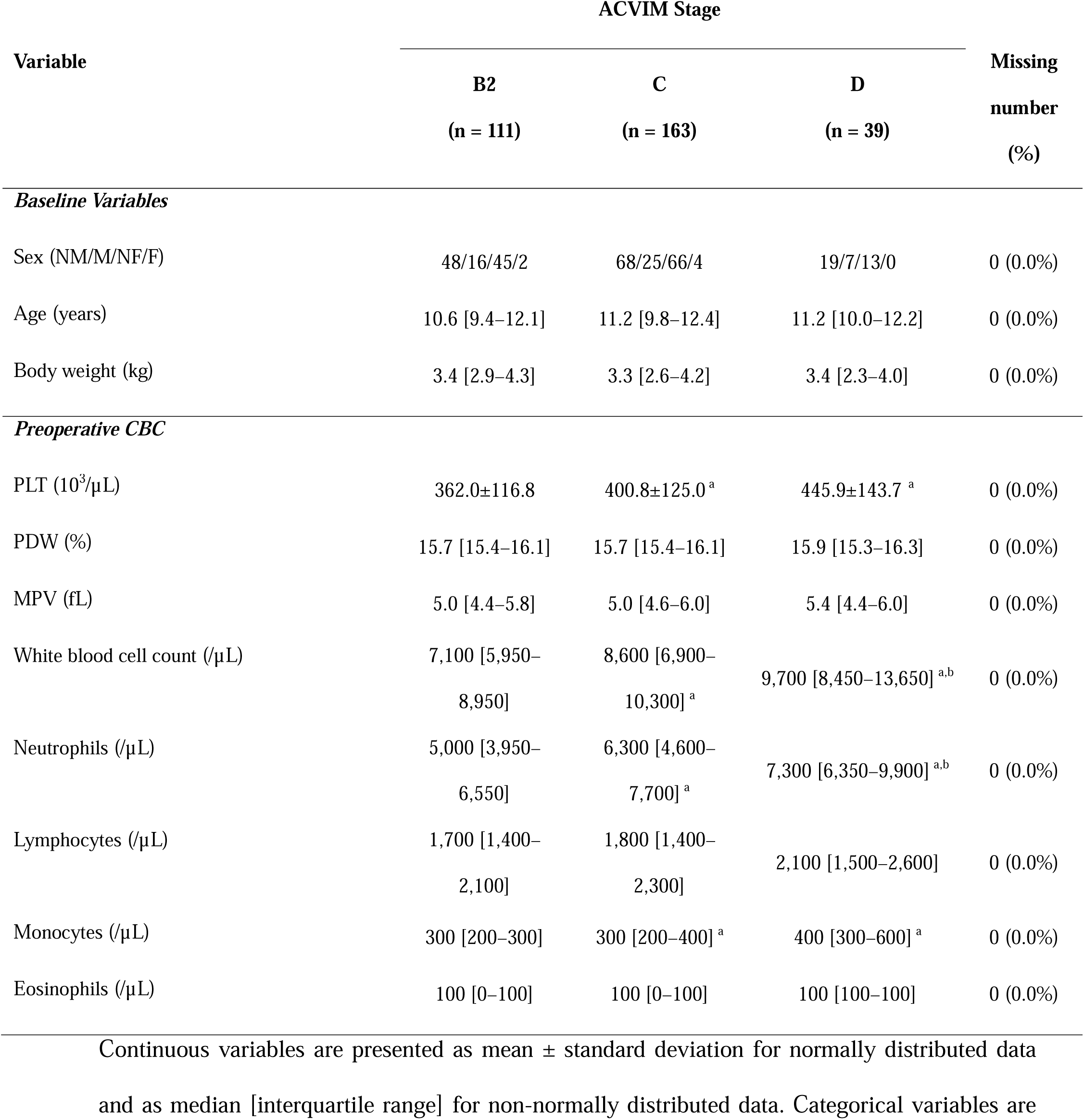
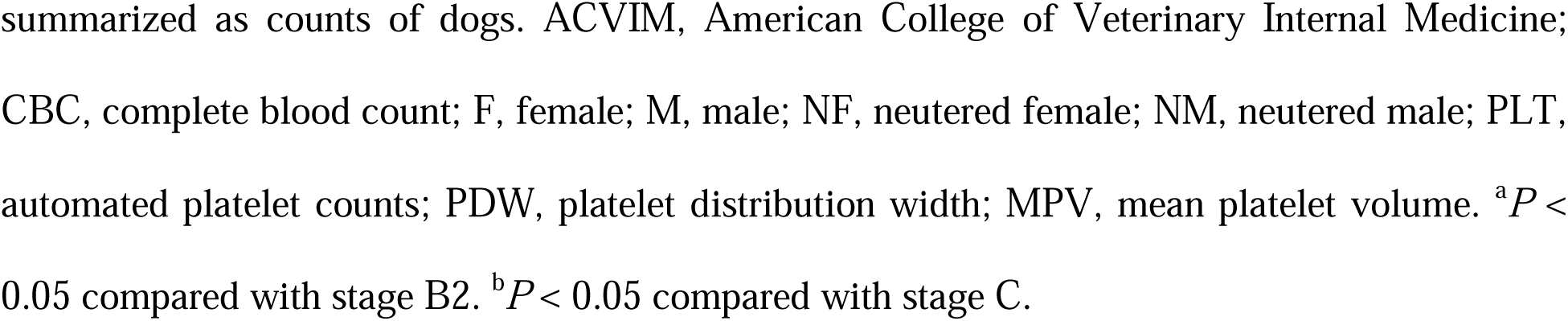
Basic Characteristics of Dogs Undergoing Mitral Valve Repair, Stratified by ACVIM Stage (B2, C, and D)

| Variable | ACVIM Stage |  |  | Missing<br>number<br>(%) |
| --- | --- | --- | --- | --- |
|  | B2 | C | D |  |
|  | (n = 111) | (n = 163) | (n = 39) |  |
| *Baseline Variables* |  |  |  |  |
| Sex (NM/M/NF/F) | 48/16/45/2 | 68/25/66/4 | 19/7/13/0 | 0 (0.0%) |
| Age (years) | 10.6 [9.4–12.1] | 11.2 [9.8–12.4] | 11.2 [10.0–12.2] | 0 (0.0%) |
| Body weight (kg) | 3.4 [2.9–4.3] | 3.3 [2.6–4.2] | 3.4 [2.3–4.0] | 0 (0.0%) |
| *Preoperative CBC* |  |  |  |  |
| PLT (10<sup>3</sup>/μL) | 362.0±116.8 | 400.8±125.0 <sup>a</sup> | 445.9±143.7 <sup>a</sup> | 0 (0.0%) |
| PDW (%) | 15.7 [15.4–16.1] | 15.7 [15.4–16.1] | 15.9 [15.3–16.3] | 0 (0.0%) |
| MPV (fL) | 5.0 [4.4–5.8] | 5.0 [4.6–6.0] | 5.4 [4.4–6.0] | 0 (0.0%) |
| White blood cell count (/μL) | 7,100 [5,950–<br>8,950] | 8,600 [6,900–<br>10,300] <sup>a</sup> | 9,700 [8,450–13,650] <sup>a,b</sup> | 0 (0.0%) |
| Neutrophils (/μL) | 5,000 [3,950–<br>6,550] | 6,300 [4,600–<br>7,700] <sup>a</sup> | 7,300 [6,350–9,900] <sup>a,b</sup> | 0 (0.0%) |
| Lymphocytes (/μL) | 1,700 [1,400–<br>2,100] | 1,800 [1,400–<br>2,300] | 2,100 [1,500–2,600] | 0 (0.0%) |
| Monocytes (/μL) | 300 [200–300] | 300 [200–400] <sup>a</sup> | 400 [300–600] <sup>a</sup> | 0 (0.0%) |
| Eosinophils (/μL) | 100 [0–100] | 100 [0–100] | 100 [100–100] | 0 (0.0%) |

Continuous variables are presented as mean ± standard deviation for normally distributed data and as median [interquartile range] for non-normally distributed data. Categorical variables are
summarized as counts of dogs. ACVIM, American College of Veterinary Internal Medicine; CBC, complete blood count; F, female; M, male; NF, neutered female; NM, neutered male; PLT, automated platelet counts; PDW, platelet distribution width; MPV, mean platelet volume. <sup>a</sup>*P* < 0.05 compared with stage B2. <sup>b</sup>*P* < 0.05 compared with stage C.

Regarding intraoperative findings, the median intubation time was 179 min [159–197 min], aortic cross-clamp time was 57 min [46–66 min], and total CPB time was 77 min [65–89 min]. Autotransfusion was used in 112 of 313 dogs, and intraoperative donor transfusion was used in 80 of 313 dogs. Among donor transfused dogs, 76 received packed red blood cells only, three received both packed red blood cells and fresh frozen plasma, and one received fresh frozen plasma alone. Median cardioplegia volume was 19.5 mL/kg [16.9–24.0 mL/kg], and intraoperative re-clamping was required in 11 of 313 dogs (**Table 2**). Eight dogs died within 5 days postoperatively, and their PLT values remained in the analysis.

**Table 2.** Intraoperative Findings of Dogs Undergoing Mitral Valve Repair, Stratified by ACVIM Stage (B2, C, and D)

| Variable | ACVIM Stage |  |  | Missing<br>number<br>(%) |
| --- | --- | --- | --- | --- |
|  | B2<br>(n = 111) | C<br>(n = 163) | D<br>(n = 39) |  |
| Time intubated (min) | 173 [158–188] | 180 [159–202] | 183 [167–196] | 36 (11.5%) |
| Cross-clamp time (min) | 53 [44–64] | 58 [48–67] | 60 [49–68] | 0 (0.0%) |
| Total CPB time (min) | 74 [64–86] | 79 [67–90] | 80 [70–95] | 0 (0.0%) |
| Cardioplegia solution volume (mL/kg) | 18.4 [16.1–22.1] | 20.2 [17.2–24.1] | 22.5 [28.7–27.7]<sup>a,b</sup> | 0 (0.0%) |
| Intraoperative autotransfusion (Yes/No) | 77/34 | 100/63 | 24/15 | 0 (0.0%) |
| Intraoperative donor transfusion (Yes/No) | 27/84 | 42/121 | 11/28 | 0 (0.0%) |
| Intraoperative re-clamp (Yes /No) | 4/107 | 5/158 | 2/37 | 0 (0.0%) |

Continuous variables are presented as median [interquartile range]. Categorical variables are summarized as counts of dogs. ACVIM, American College of Veterinary Internal Medicine; CBP, cardiopulmonary bypass. <sup>a</sup>*P* < 0.05 compared with stage B2. <sup>b</sup>*P* < 0.05 compared with stage C.

Mean PLT values were 393±127 ×10³/µL on Day 0, 159±72 on Day 1, 137±67 on Day 2, 157±87 on Day 3, and 187±104 on Day 4. Postoperative thrombocytopenia during Days 1–4 occurred in 124 of 313 dogs (**Figure 1**). In the multivariable mixed effects model, PLT (unit: ×10³/µL) counts decreased markedly from Day 0 to Day 1 (β = −207.33 [×10³/µL per day], 95% CI: −290.98 to −123.67, P < 0.001) and showed a recovery trend from Day 1 to 4 (β = 20.62 [×10³/µL per day], 95% CI: −2.35 to 43.60, P = 0.079). PLT across the observation period were associated with age, breed (Toy Poodle and Yorkshire Terrier), ACVIM stage, neuter status, and body weight. Older dogs had higher PLT (β = 17.39 [×10³/µL per year], 95% CI: 9.68 to 25.11, P < 0.001). Toy Poodles (β = −107.68 [×10³/µL], 95% CI: –145.35 to –70.01, P < 0.001) and Yorkshire Terriers (β = −129.36 [×10³/µL], 95% CI: –216.31 to –42.41, P = 0.004) had lower PLT than Chihuahuas. Dogs in ACVIM stages C and D had higher PLT than those in B2 (Stage C: β = 27.43 [×10³/µL], 95% CI: 0.05–54.82, P = 0.050; Stage D: β = 88.65 [×10³/µL], 95% CI: 46.13–131.18, P < 0.001). Lower body weight was associated with lower PLT (β = −14.95 [×10³/µL per kg], 95% CI: –28.81 to –1.09, P = 0.035). Intact males (β = 37.49 [×10³/µL], 95% CI: 0.13–74.84, P = 0.049) and intact females (β = 127.14 [×10³/µL], 95% CI: 35.04–219.23, P = 0.007) had higher PLT than neutered males (**Figure 2**). Covariates associated with higher preoperative PLT tended to show greater decreases from Day 0 to 1. Toy Poodles demonstrated less recovery from Day 1 to 4 compared with Chihuahuas (β = −10.67 [×10³/µL per day]; 95% CI: –18.98 to –2.37; P = 0.012) (**Supplemental Table 1**). PDW increased from Day 0 to 1 and remained stable thereafter. MPV was stable from Day 0 to 1 and showed a mild increasing trend from Day 1 to 4 (**Supplemental Figure 1**). Shih Tzus and Pomeranians had higher PDW values, and Toy Poodles had higher MPV values than Chihuahuas (**Supplemental Figure 2**).

**Figure 1.**
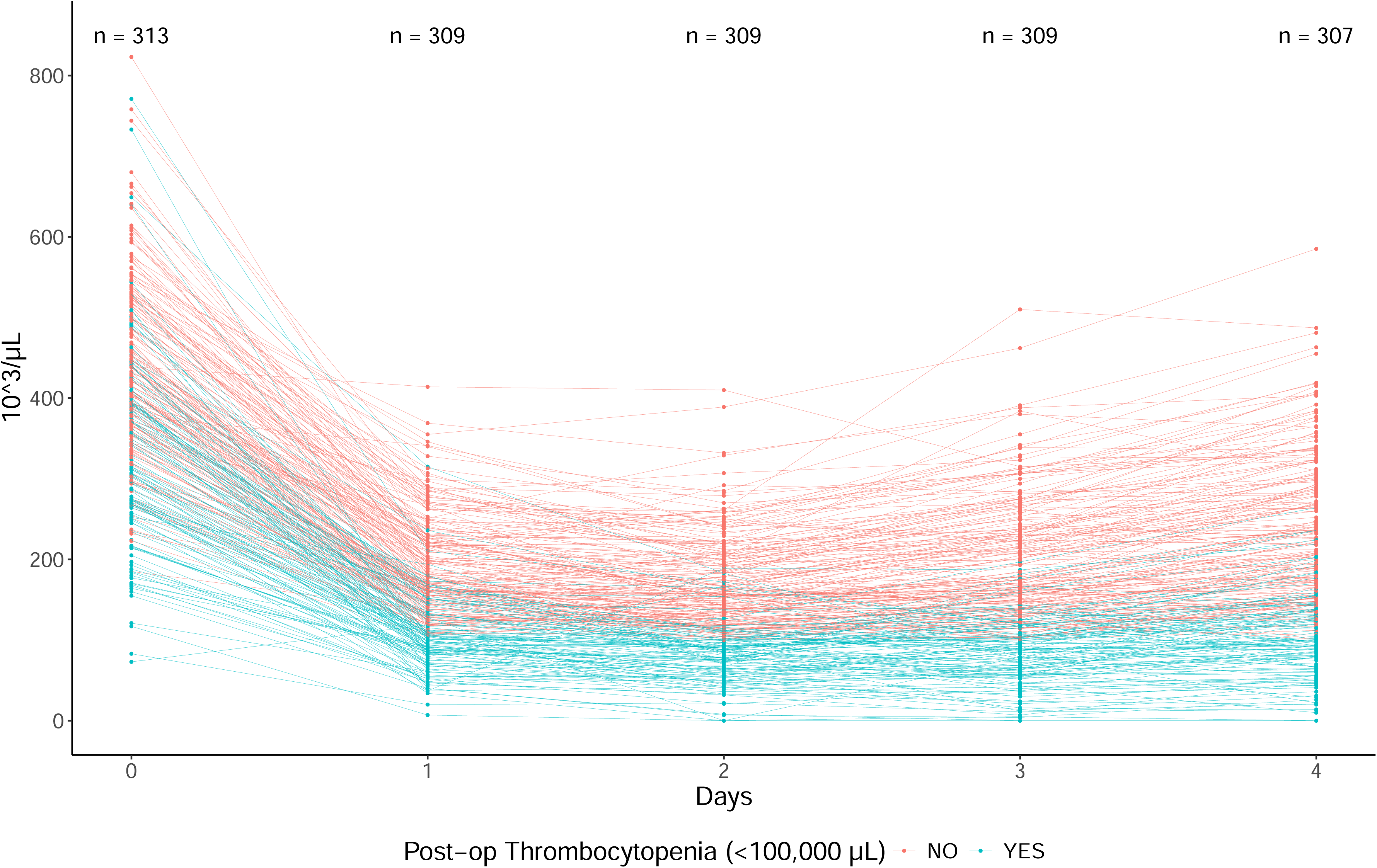
Spaghetti Plot of Automated Platelet Count Trajectories in Dogs With and Without Postoperative Thrombocytopenia. Spaghetti plot showing individual platelet count trajectories from Day 0 to Day 4, with dogs experiencing postoperative thrombocytopenia (platelet count <100 ×10³/µL) between Day 1 and Day 4 shown in blue.

**Figure 2.**
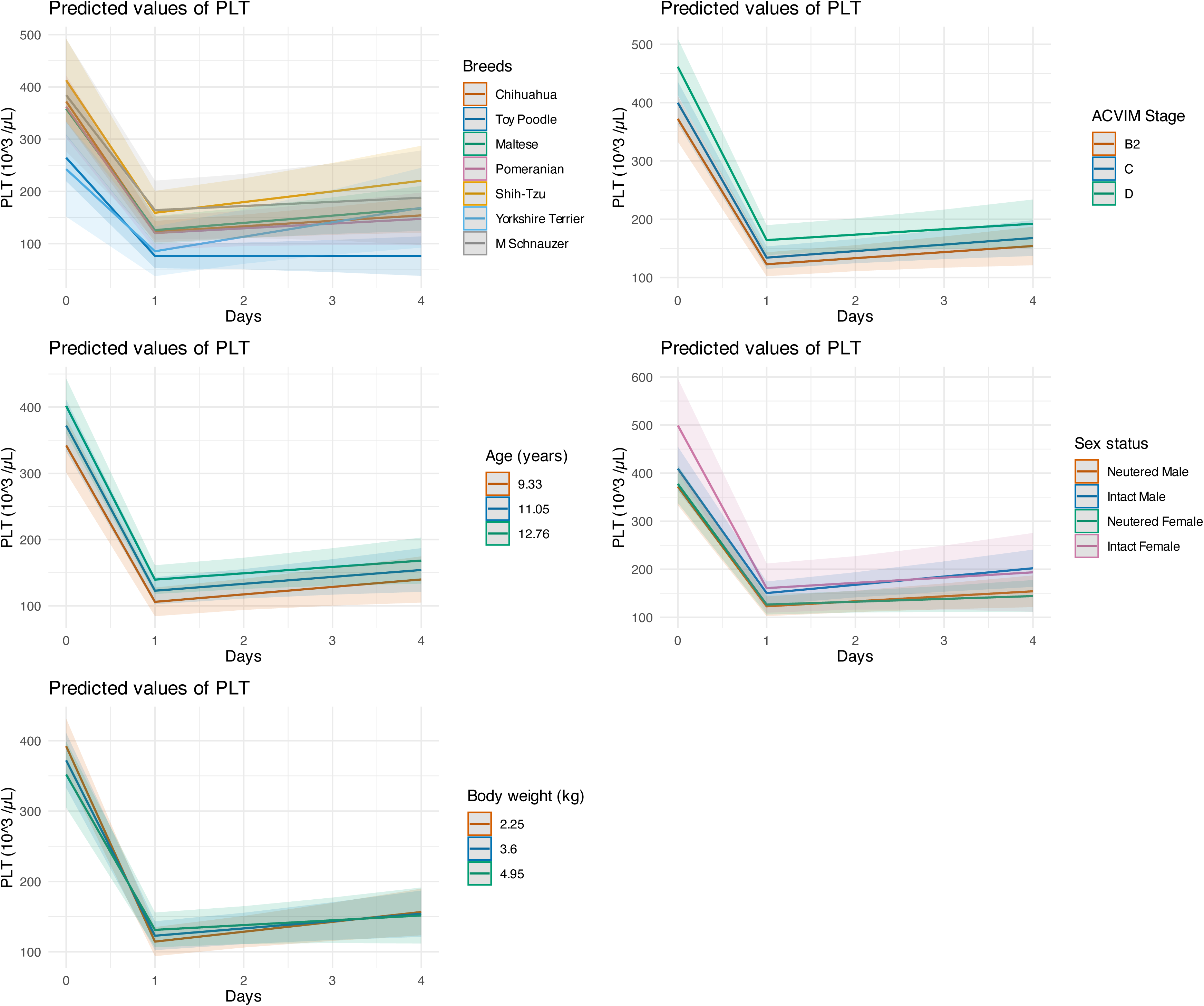
Predicted Platelet Count Trajectories Based on Mixed-Effects Model Results. Predicted platelet count trajectories from the multivariable mixed-effects model with linear splines (knot at Day 1) are shown across key clinical variables: (A) breed, (B) ACVIM stage, (C) age, (D) sex status, and (D) body weight. Each plot presents predicted means with 95% confidence intervals derived from the final model (Table 2). The model includes time as a fixed effect that interacts with clinical variables and as a random effect to account for variation in slopes and intercepts among individual dogs. For age and body weight, modeled as continuous variables, representative values such as the lower quartile, median, and upper quartile were selected to support clinical interpretation.

Postoperatively, 18 dogs received transfusions for bleeding or declining Hct levels: two received whole blood, one received packed red blood cells, eight received fresh frozen plasma (FFP), and seven received both FFP and packed red blood cells. Most of (14 out of 18) these transfusions occurred immediately after surgery, before the CBC recheck on Day 1. These dogs showed a visually notable steep decline in PLT from Day 0 to 1 (**Supplemental Figure 3**).

## Discussions

The results showed that mean PLT counts declined immediately after surgery and gradually recovered from Day 1 to Day 4. Toy Poodles and Yorkshire Terriers had lower PLT levels than Chihuahuas, whereas intact males and intact females had higher PLT than neutered males. Dogs in advanced ACVIM stages also tended to have higher PLT. Postoperative thrombocytopenia occurred in 124 of 313 dogs, demonstrating that it was a frequent finding following MVR. This study was the first to describe postoperative PLT trends in dogs undergoing MVR, providing important information for future cohort studies and postoperative management in veterinary practice.

Cardiopulmonary bypass is known to cause blood trauma, including red blood cell destruction and PLT activation from shear stress [16]. Hemodilution further affects blood cell concentrations, especially in small patients. At our institution, the CPB priming volume is at least 140 mL, while the median body weight of the population is 3.3 kg, producing a dilution ratio of approximately 0.68. Even with packed red blood cell transfusions, PLT-related problems remain uncorrected. This is a significant challenge in veterinary medicine, where PLT transfusion is not routinely available. In humans, thrombocytopenia within 4 postoperative days is largely attributed to hemodilution and perioperative PLT consumption before thrombopoietin-driven recovery and transient overshoot [8]. Most human cardiac surgery patients reach a PLT nadir on postoperative Days 2–3, returning to baseline by Day 5 [17, 18], and 97% of human patients in a prospective cohort of 581 reached PLT nadir between Day 1 and 4 [19, 20]. The current study demonstrated a transient decrease in PLT on Day 1 and gradual increase through Day 4.

In studies of older individuals, PLT progressively increased with age, likely due to increased inflammatory burden [21]. In the present study, higher age was also associated with higher PLT across the observation period; however, the interaction between age and time was minimal, indicating that the magnitude of postoperative decline and the rate of recovery did not meaningfully vary with age.

Although this study excluded breeds predisposed to inherent PLT disorders, Toy Poodles and Yorkshire Terriers still exhibited lower PLT levels than Chihuahuas. Prior work reported that Yorkshire Terriers have a breed-specific reference interval lower than standard laboratory ranges [22] and another study showed Poodles often have lower PLT values than Beagles, Maltese, and Schnauzers [23]. These findings suggest that baseline breed-related differences in PLT should be considered when interpreting postoperative PLT levels. Notably, Toy Poodle was the only variable that influenced PLT recovery during Days 1–4, showing a slower rise compared with Chihuahuas. Whether this phenomenon reflected delayed PLT regeneration or resulted from breed-related variability in platelet size that affects automated PLT counting remains unclear. Further studies incorporating manual PLT counts in predisposed breeds are warranted to clarify differences in postoperative recovery patterns.

As previously reported, PLT tended to increase with progression of MMVD severity in dogs [24, 25]. Platelet release pro-inflammatory mediators, and activated platelets play a key role in cardiovascular disease by promoting thrombus formation after endothelial injury [26, 27]. Megakaryocytes, the precursor cells for platelets, are strongly influenced by inflammation; cytokines such as interleukin-6 and interleukin-1 enhance platelet production by promoting megakaryocyte maturation and increasing thrombopoietin synthesis [28]. Shear stress caused by mitral regurgitation may also alter platelet function and shorten platelet lifespan in dogs [25]. In the present study, as in prior reports, higher preoperative PLT were observed in dogs with advanced ACVIM stages and remained higher throughout the postoperative period, although recovery patterns did not differ among stages. Higher baseline neutrophil counts were also observed in advanced stages, suggesting that sustained inflammation may contribute to elevated PLT.

Intact sex status appeared to be associated with higher PLT during observation period. Intact females had the highest preoperative PLT, followed by intact males, with both neutered groups showing lower counts. These trends also persisted postoperatively, although the recovery trajectory did not differ significantly among sex status. Increased PLT in female dogs has been reported previously [29, 30], and the interaction between sex and neutering status has also been shown to influence PLT [22]. Similar patterns have been noted in species as diverse as rock hyraxes [31] and humans [21], suggesting that sex hormones exert conserved effects on PLT dynamics across evolutionarily distant species. Androgens may promote myeloid differentiation [32, 33], whereas estrogens may favor megakaryocytic differentiation [34].

Platelet consumption and hemodilution are considered the primary causes of PLT decline after cardiac surgery with CPB. Once CPB began, platelets were activated by mechanical shear stress and exposure to artificial surfaces, initiating an activation cascade [35]. Such activation has been shown to lead to a preferential loss of larger and more reactive platelets, presumably due to blood–membrane contact and shear forces within the extracorporeal circuit, resulting in their subsequent sequestration and consumption in the peripheral circulation [36]. High shear stress may also induce platelet apoptosis [37]. In human studies, prolonged CPB time has been identified as a risk factor for postoperative thrombocytopenia, with patients developing thrombocytopenia exhibiting CPB durations of 173 ± 79 min compared with 131 ± 60 min in those without thrombocytopenia [38]. In contrast, the present study found no significant association between PLT and CPB time. This difference may reflect the uniform surgical technique and relatively narrow range of CPB durations in the cohort, with a median CPB time of 77 min [65–89 min] and no cases requiring markedly prolonged bypass. Future studies including more complex procedures or broader case diversity may clarify whether CPB duration affects PLT dynamics.

Regarding intraoperative transfusion, autotransfusion did not affect PLT. This technique was used to reduce hemodilution and blood cell injury during CPB and to limit loss of coagulation factors; however, it did not demonstrate a protective effect on PLT counts. Intraoperative donor transfusion with packed red blood cells was administered when the expected Hct after priming dilution fell below 20%. Although intended to maintain Hct levels, this approach did not offset platelet loss. Therefore, the persistent postoperative decrease in PLT likely reflected unavoidable hemodilution despite transfusion strategies.

Postoperative factors, such as the amount of chest tube drainage (as an indicator of bleeding), the presence of purpura, postoperative transfusion use, and the effects of anticoagulants, were not included as covariates because of the retrospective design. Postoperative bleeding and purpura could contribute to thrombocytopenia through platelet loss and consumption. Whole blood transfusion may directly increase PLT, while FFP may influence counts indirectly by supplying coagulation factors. In this study, two dogs received whole blood and fifteen received FFP during Days 1–4. To better evaluate relationships between transfusion and PLT, future investigations incorporating coagulation factor changes and transfusion timing as time-dependent covariates are needed.

Regarding anticoagulants, dalteparin (or rivaroxaban) and clopidogrel were routinely administered during hospitalization according to institutional protocol. Thrombotic thrombocytopenic purpura has been reported in humans after initiation of clopidogrel, typically within the first 2 weeks [39]. Drug-induced immune thrombocytopenia, particularly heparin-induced thrombocytopenia, may lead to hemorrhage and generally occurs 5–10 days after exposure [40]. Because this study focused solely on early postoperative PLT trends, these medications were unlikely to have directly influenced the PLT decline. However, the combination of anticoagulant use and decreased coagulation factors may have promoted bleeding tendencies, including hemorrhage or purpura, which could have contributed to subsequent reductions in PLT. Longer follow-up is warranted to assess possible adverse drug effects.

This study has several limitations. First, although routine examinations, including CBC, were performed daily until Day 4, the dataset lacked sufficient information to assess hemostatic changes beyond 5 days postoperatively. A typical biphasic thrombocytopenia pattern, consisting of an initial postoperative decline, recovery, and a delayed secondary drop potentially related to immune-mediated mechanisms such as heparin-induced thrombocytopenia, could not be assessed. Second, the precision of statistical estimates may have been limited by model assumptions, small subgroup sizes, and unmeasured confounders, including comorbidities affecting PLT. Third, the effects of perioperative anticoagulants and potential measurement errors (differences in personnel, sampling technique, blood collection sites, and automated CBC accuracy) could not be controlled due to the retrospective design. The absence of manual PLT data creates uncertainty regarding whether low PLT values reflected true thrombocytopenia. Missing PLT data from dogs that died in hospital may also introduce survival bias. Finally, external validity may be restricted, as the study excluded Cavalier King Charles Spaniels, a major breed undergoing MVR in England [41, 42], limiting generalizability to broader breed populations.

In conclusion, a marked decrease in PLT occurred after MVR, followed by a mild increasing trend between Day 1 and 4. PLT dynamics were associated with breed, ACVIM stage, and sex status. Recognizing the relationships between PLT changes and baseline variables may help clinicians identify high-risk patients and apply targeted strategies, such as enhanced monitoring, premedication, or transfusion, to optimize outcomes for dogs with MMVD undergoing MVR.

## Abbreviations

ACVIM: American College of Veterinary Internal Medicine
CBC: Complete blood count
CI: Confidence interval
CPB: Cardiopulmonary bypass
HR: Heart rate
MVR: Mitral valve repair
MMVD: Myxomatous mitral valve disease
PLT: Platelet count
RR: Risk ratio

---

**Supplemental Table 1.** Mixed-Effects Model Results for Perioperative Platelet Count (×10^3^/µL) Changes in Dogs Undergoing Mitral Valve Repair.

| Variable | Estimate | Lower 95% CI | Upper 95% CI | P-value |
| --- | --- | --- | --- | --- |
| ***Main Effect*** |  |  |  |  |
| Day 0–1 | -207.33 | -290.98 | -123.67 | <0.001 |
| Day 1–4 | 20.62 | -2.35 | 43.60 | 0.079 |
| Age (year) | 17.39 | 9.68 | 25.11 | <0.001 |
| Toy Poodle (vs. Chihuahua) | -107.68 | -145.35 | -70.01 | <0.001 |
| Maltese (vs. Chihuahua) | -13.96 | -55.07 | 27.16 | 0.506 |
| Pomeranian (vs. Chihuahua) | -10.16 | -61.20 | 40.88 | 0.697 |
| Shih Tzu (vs. Chihuahua) | 38.29 | -40.29 | 116.87 | 0.340 |
| Yorkshire Terrier (vs. Chihuahua) | -129.36 | -216.31 | -42.41 | 0.004 |
| Miniature Schnauzer (vs. Chihuahua) | 12.06 | -93.33 | 117.46 | 0.823 |
| ACVIM stage C (vs. B2) | 27.43 | 0.05 | 54.82 | 0.050 |
| ACVIM stage D (vs. B2) | 88.65 | 46.13 | 131.18 | <0.001 |
| Body weight (kg) | -14.95 | -28.81 | -1.09 | 0.035 |
| Intact male (vs. Neutered male) | 37.49 | 0.13 | 74.84 | 0.049 |
| Neutered female (vs. Neutered male) | 5.36 | -22.77 | 33.49 | 0.709 |
| Intact female (vs. Neutered male) | 127.14 | 35.04 | 219.23 | 0.007 |
| Total CPB time (per 10 min) | 2.68 | -4.69 | 10.04 | 0.476 |
| Cardioplegia solution volume (mL/kg) | -1.77 | -4.11 | 0.57 | 0.138 |
| Intraoperative autotransfusion (Yes vs. No) | 7.95 | -23.16 | 39.05 | 0.616 |
| Intraoperative donor transfusion (Yes vs. No) | 17.12 | -16.14 | 50.38 | 0.313 |
| Intraoperative re-clamp (Yes vs. No) | -38.19 | -112.21 | 35.82 | 0.312 |
| ***Interaction with time<sup>†</sup>*** |  |  |  |  |
| Age (year) × Day 0–1 | -7.56 | -13.71 | -1.41 | 0.016 |
| Age (year) × Day 1–4 | -0.47 | -2.18 | 1.23 | 0.585 |
| Toy Poodle × Day 0–1 | 61.02 | 30.96 | 91.09 | 0.000 |
| Toy Poodle × Day 1–4 | -10.67 | -18.98 | -2.37 | 0.012 |
| Maltese × Day 0–1 | 17.00 | -15.73 | 49.73 | 0.309 |
| Maltese × Day 1–4 | 3.40 | -5.59 | 12.39 | 0.459 |
| Pomeranian × Day 0–1 | 4.63 | -35.99 | 45.26 | 0.823 |
| Pomeranian × Day 1–4 | -1.79 | -12.97 | 9.39 | 0.754 |
| Shih Tzu × Day 0–1 | 0.03 | -62.53 | 62.58 | 0.999 |
| Shih Tzu × Day 1–4 | 10.23 | -7.03 | 27.49 | 0.245 |
| Yorkshire Terrier × Day 0–1 | 91.71 | 22.51 | 160.91 | 0.009 |
| Yorkshire Terrier × Day 1–4 | 17.42 | -1.54 | 36.39 | 0.072 |
| Miniature Schnauzer × Day 0–1 | 27.69 | -56.19 | 111.58 | 0.518 |
| Miniature Schnauzer × Day 1–4 | -2.74 | -25.77 | 20.29 | 0.816 |
| ACVIM stage C × Day 0–1 | -16.40 | -38.21 | 5.41 | 0.141 |
| ACVIM stage C × Day 1–4 | 0.81 | -5.20 | 6.81 | 0.793 |
| ACVIM stage D × Day 0–1 | -46.40 | -80.38 | -12.42 | 0.007 |
| ACVIM stage D × Day 1–4 | -1.02 | -10.60 | 8.55 | 0.834 |
| Body weight (kg) × Day 0–1 | 21.62 | 10.59 | 32.66 | 0.000 |
| Body weight (kg) × Day 1–4 | -2.55 | -5.59 | 0.50 | 0.101 |
| Intact male × Day 0–1 | -9.67 | -39.50 | 20.15 | 0.525 |
| Intact male × Day 1–4 | 6.76 | -1.48 | 15.00 | 0.108 |
| Neutered female × Day 0–1 | -1.22 | -23.63 | 21.19 | 0.915 |
| Neutered female × Day 1–4 | -4.45 | -10.61 | 1.70 | 0.156 |
| Intact female × Day 0–1 | -88.94 | -162.23 | -15.64 | 0.017 |
| Intact female × Day 1–4 | 0.56 | -19.50 | 20.62 | 0.956 |
| Total CPB time (per 10 min) × Day 0–1 | -5.83 | -11.70 | 0.04 | 0.051 |
| Total CPB time (per 10 min) × Day 1–4 | -0.22 | -1.88 | 1.44 | 0.797 |
| Cardioplegia solution volume (mL/kg) × Day 0–1 | 0.47 | -1.39 | 2.34 | 0.619 |
| Cardioplegia solution volume (mL/kg) × Day 1–4 | 0.28 | -0.26 | 0.82 | 0.313 |
| Intraoperative autotransfusion (Yes) × Day 0–1 | 14.76 | -10.03 | 39.55 | 0.243 |
| Intraoperative autotransfusion (Yes) × Day 1–4 | 0.84 | -6.00 | 7.67 | 0.811 |
| Intraoperative donor transfusion (Yes) × Day 0–1 | -27.82 | -54.41 | -1.23 | 0.040 |
| Intraoperative donor transfusion (Yes) × Day 1–4 | 3.78 | -3.61 | 11.16 | 0.316 |
| Intraoperative re-clamp (Yes) × Day 0–1 | 19.08 | -39.85 | 78.01 | 0.526 |
| Intraoperative re-clamp (Yes) × Day 1–4 | -1.29 | -17.65 | 15.08 | 0.878 |

†The interaction term reflects differences in how PLT levels change over time between groups
defined by each covariate in main effects.

**Supplemental Figure 1. Spaghetti Plot of Platelet Distribution Width (PDW) and Mean Platelet Volume (MPV) Trajectories in Dogs With and Without Postoperative Thrombocytopenia.**

Spaghetti plot showing individual PDW (A) and MPV (B) trajectories from Day 0 to Day 4. Dogs with postoperative thrombocytopenia (platelet count <100 × 10³/µL) between Day 1 and Day 4 are depicted in blue.

**Supplemental Figure 2. Predicted Platelet Distribution Width (PDW) and Mean Platelet Volume (MPV) Trajectories Based on Mixed-Effects Model Results.**

Predicted PDW (A) and MPV (B) from the multivariable mixed-effects model with linear splines (knot at Day 1) are displayed across breeds. Each plot presents predicted means with 95% confidence intervals. The model incorporates time as a fixed effect that interacts with clinical variables and as a random effect allowing variation in slopes and intercepts among dogs. PDW was higher in Pomeranian (β = 0.4607 [%]; 95% CI: 0.1313–0.7902; P = 0.006) and Shih Tzu (β = 0.6797 [%]; 95% CI: 0.1724–1.1869; P = 0.009) compared with Chihuahua. MPV was higher in Toy Poodle (β = 0.9480 [fL]; 95% CI: 0.5977–1.2984; P < 0.001) compared with Chihuahua.

**Supplemental Figure 3. Spaghetti Plot of Automated Platelet Count Trajectories in Dogs With and Without Postoperative Transfusion.**

Spaghetti plot showing individual platelet count trajectories from Day 0 to Day 4. Dogs with postoperative transfusion between Day 1 and Day 4 (n = 18) are depicted in red: two received whole blood, one received packed red blood cells, eight received fresh frozen plasma (FFP), and seven received both FFP and packed red blood cells. Most (14 of 18) were transfused immediately after surgery, before the CBC recheck on Day 1.

## Notes

### Competing Interest Statement

The authors have declared no competing interest.

