## Supplementary figures and images for "Postoperative Automated Platelet Count and Its Predictive Factors in Dogs Undergoing Mitral Valve Repair"

### Supplemental Figure 1

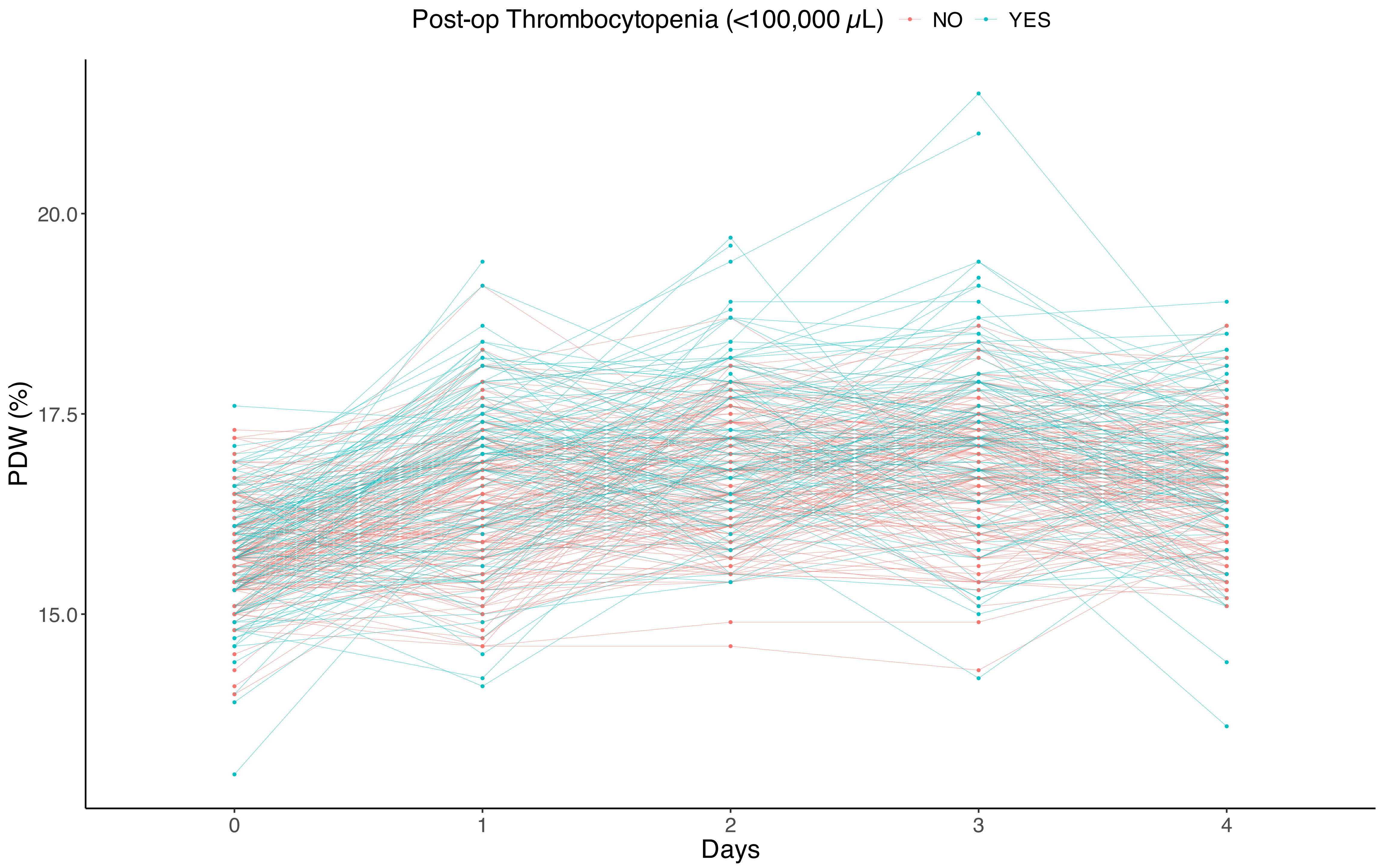

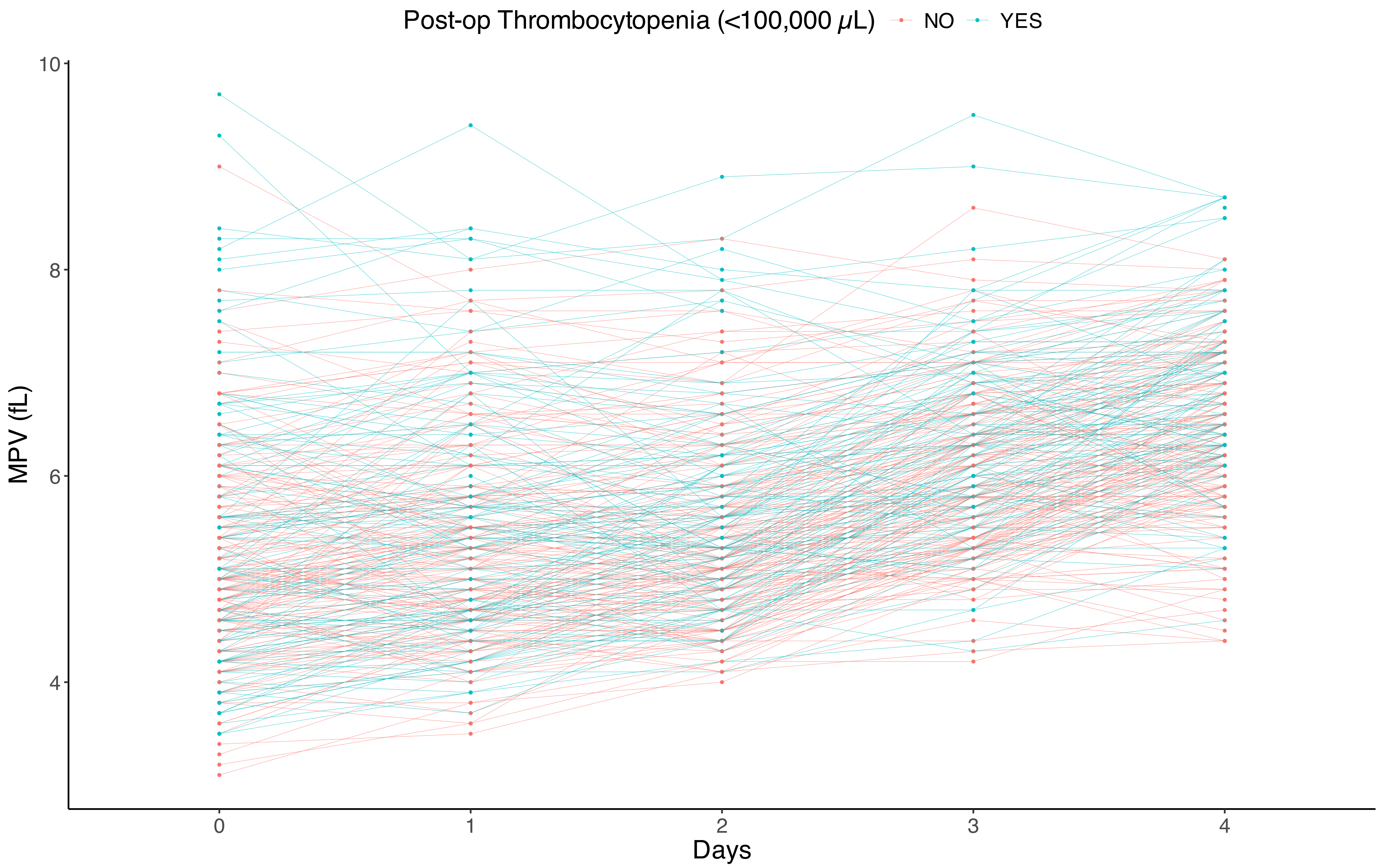

### Supplemental Figure 2

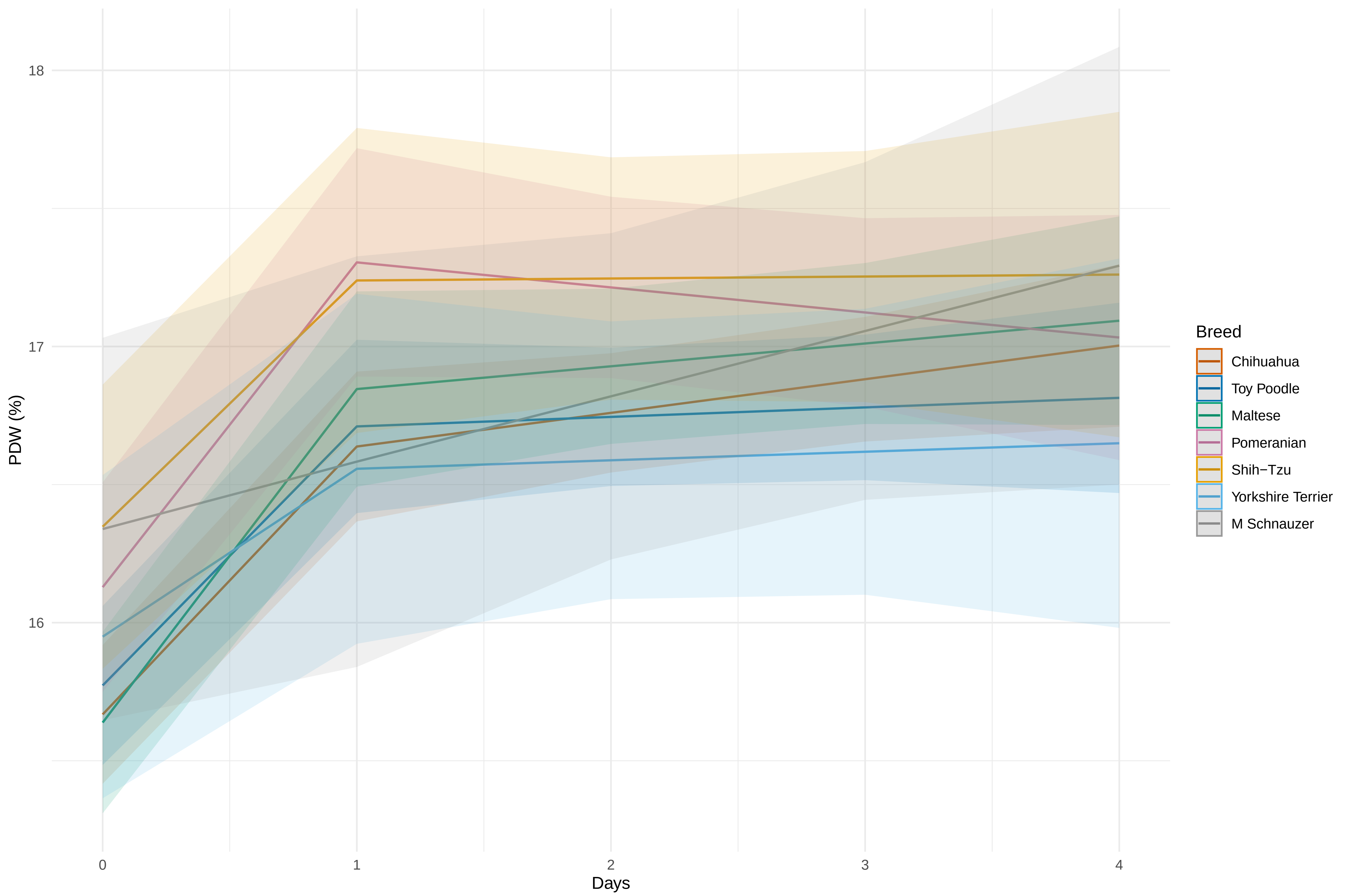

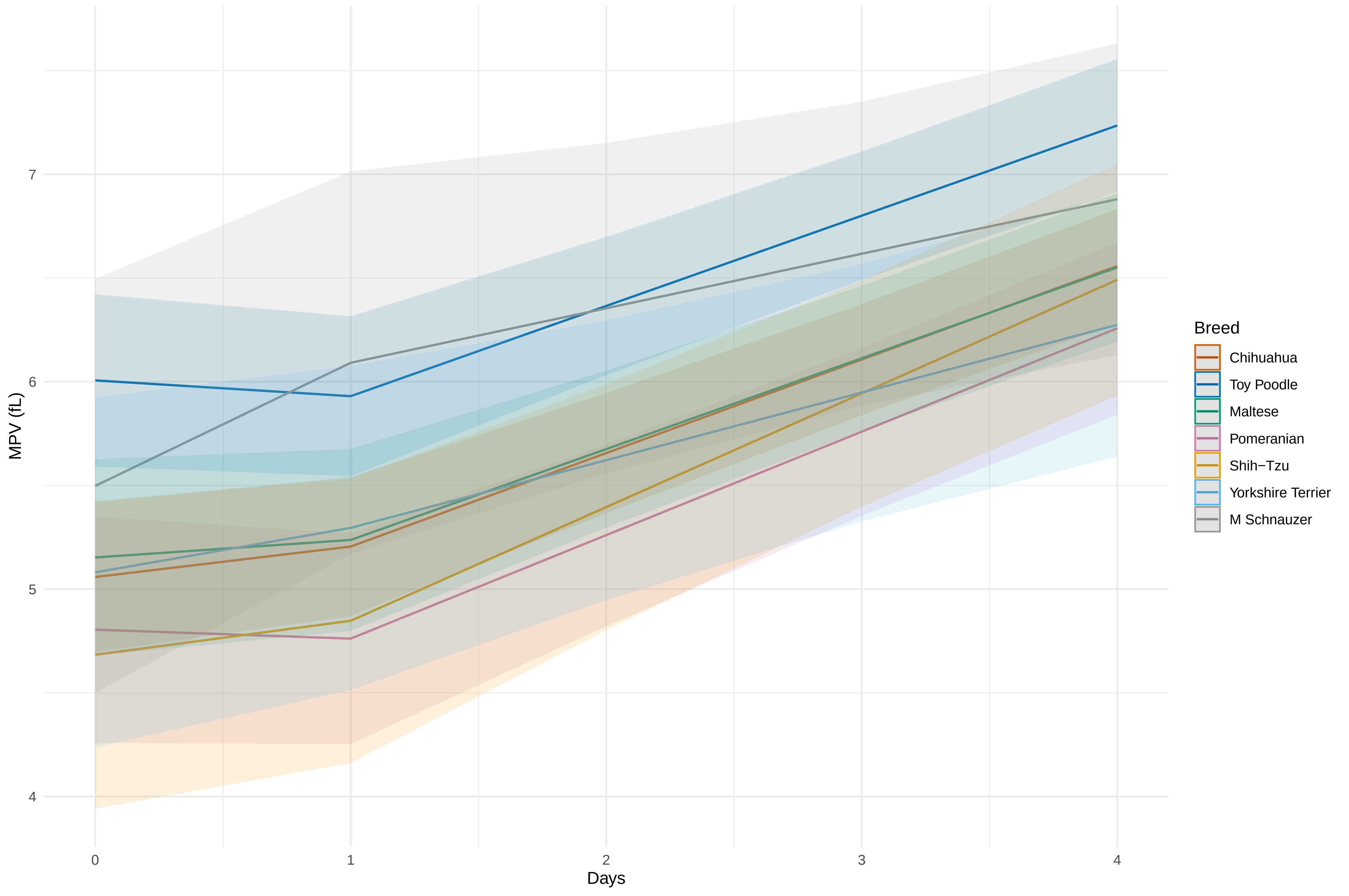

### Supplemental Figure 3

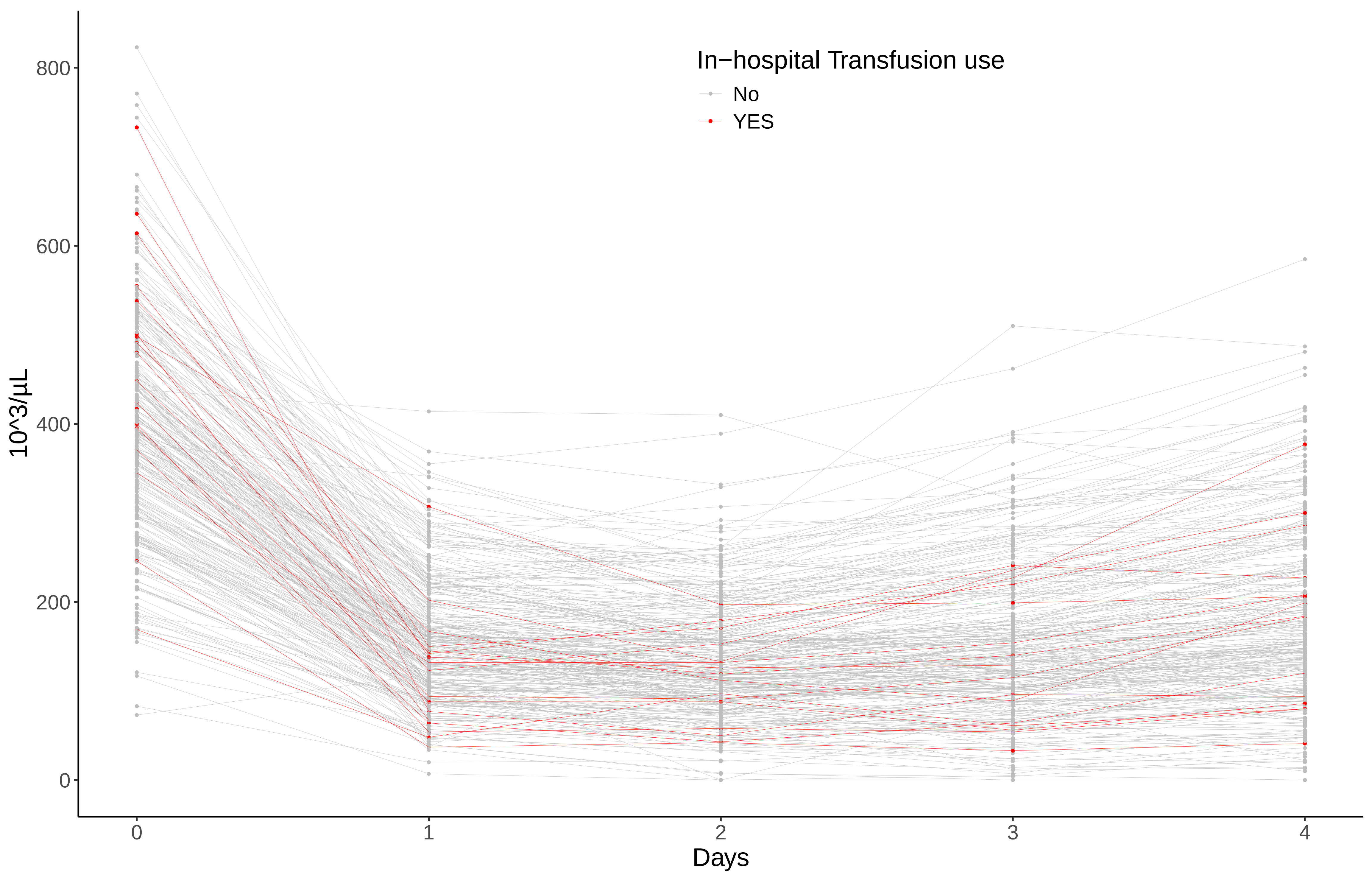
